# Chronic Alcohol Dysregulates Glutamatergic Function in the Basolateral Amygdala in a Projection- and Sex-Specific Manner

**DOI:** 10.1101/2021.12.22.473716

**Authors:** Michaela E. Price, Brian A. McCool

## Abstract

Chronic intermittent ethanol (CIE) produces alcohol dependence, facilitates anxiety-like behavior, and increases post-CIE alcohol intake. The basolateral amygdala (BLA) is one of several brain regions that regulates anxiety-like behavior and alcohol intake through downstream projections. The BLA receives information from two distinct input pathways. Afferents from medial structures like the thalamus and prefrontal cortex enter the BLA through the *stria terminalis* whereas lateral cortical structures like the anterior insula cortex enter the BLA through the external capsule. CIE induces input- and sex-specific adaptations to glutamatergic function in the BLA. Previous studies sampled neurons throughout the BLA, but did not distinguish between projection-specific populations. The current study investigated BLA neurons that project to the NAC (BLA-NAC neurons) or the BNST (BLA-BNST neurons) as representative ‘reward’ and ‘aversion’ BLA neurons, and showed that CIE alters glutamatergic function and excitability in a projection- and sex-specific manner. CIE increases glutamate release from *stria terminalis* inputs only onto BLA-BNST neurons. At external capsule synapses, CIE increases postsynaptic glutamatergic function in male BLA-NAC neurons and female BLA-BNST neurons. Subsequent experiments demonstrated that CIE enhanced the excitability of male BLA-NAC neurons and BLA-BNST neurons in both sexes when glutamatergic but not GABAergic function was intact. Thus, CIE-mediated increased glutamatergic function facilitates hyperexcitability in male BLA-NAC neurons and BLA-BNST neurons of both sexes.

## INTRODUCTION

Chronic alcohol consumption alters reward and aversion systems in the brain, initiating a cycle of excessive alcohol consumption, abstinence/withdrawal symptoms, and relapse. These effects have been modeled by exposing rodents to chronic intermittent ethanol (CIE) via vapor inhalation. Our laboratory and others have found that CIE exposure is sufficient to increase anxiety-like behavior and reward thresholds during withdrawal (WD), as well as increase post-CIE alcohol consumption in two-bottle choice paradigms (Koob et al., 2014; Koob & Volkow, 2016; McGinnis, Parrish, & McCool, 2020; McGinnis, Parrish, Chappell, et al., 2020; Morales et al., 2015, 2018; Schulteis et al., 1995). These behavioral changes mimic symptoms observed in alcohol-dependent humans, demonstrating that the CIE paradigm induces alcohol dependence in rodents. Importantly, men have a higher prevalence of alcohol use disorder (AUD) than women (Substance Abuse and Mental Health Services Administration, 2019), however, the vast majority of preclinical studies have found that female rodents have higher baseline alcohol consumption compared to males (Almeida et al., 1998; Amodeo et al., 2018; Morales et al., 2015; Scott et al., 2020; Vetter-O’Hagen & Spear, 2011). Moreover, only male rodents increase alcohol consumption after CIE/WD (Morales et al., 2015), suggesting that males are more sensitive to changes in reward systems. In contrast, withdrawal-induced anxiety-like behavior is independent of sex (Morales et al., 2015, 2018; Overstreet et al., 2004), and thus CIE/WD alters aversion systems in a similar fashion between male and female rats.

Our laboratory has investigated the neurophysiological mechanisms underlying AUD by exposing rats to CIE and examining the neurophysiological changes in the basolateral amygdala (BLA) during WD. The BLA has a well-established role in regulating anxiety-like behavior and alcohol consumption through its downstream projections (Diaz et al., 2011; Läck et al., 2007, 2008; McCool et al., 2014; McGinnis, Parrish, Chappell, et al., 2020). Glutamatergic pyramidal neurons comprise approximately 80% of the neurons in the BLA and drive BLA output to downstream brain regions (Sah et al., 2003). Pyramidal neurons receive glutamatergic afferents from two distinct input pathways. Midline structures like the medial prefrontal cortex and the polymodal sensory thalamus send projections through the *stria terminalis* whereas more lateralized cortical structures like the anterior insula cortex project through the external capsule (Leichnetz & Astruc, 1977; McDonald, 1998; McDonald et al., 1996; Sah et al., 2003). Our laboratory has shown that inputs from the *stria terminalis* undergo presynaptic facilitation during WD in CIE-exposed rats (Christian et al., 2013; McGinnis, Parrish, Chappell, et al., 2020; Morales et al., 2018; Sizer et al., 2021). Investigating inputs from specific midline structures revealed that CIE/WD increases glutamate release from inputs originating in the dorsomedial prefrontal cortex and decreases glutamate release from ventromedial prefrontal cortex inputs (McGinnis, Parrish, Chappell, et al., 2020). In contrast, external capsule inputs like those arising from the anterior insula cortex increase postsynaptic glutamatergic function following CIE/WD (Christian et al., 2012; McGinnis, Parrish, & McCool, 2020; Morales et al., 2018). These input-specific changes in glutamatergic transmission are observed in both sexes, but female rats require a longer alcohol exposure to induce the same effects (Morales et al., 2018).

Although a diverse set of inputs converge on BLA pyramidal neurons and pyramidal neurons exhibit input-specific neurophysiological changes following CIE/WD, a recent study suggests that different groups of BLA neurons receive the same set of inputs with a similar input weighting pattern (Huang et al., 2021). In contrast, there is little collateralization between BLA projections to different downstream brain regions, indicating that BLA projections are largely independent of one another and the vast majority of BLA neurons project to a single downstream target (Beyeler et al., 2016, 2018; Huang et al., 2021; Senn et al., 2014). Thus, all BLA neurons integrate information from the same set of diverse sources and then BLA neurons are segregated into projection-specific groups that independently regulate the BLA-mediated behaviors associated with each projection target. Given that most BLA neurons project to a single downstream target and there is little collateralization between BLA projections, we investigated how CIE/WD affects the neurophysiology of distinct projection-specific groups of BLA neurons.

The BLA regulates anxiety-like behavior and alcohol consumption through downstream projections to anxiety and reward-related brain regions. The BLA projection to the nucleus accumbens (NAC) is preferentially responsive to reward-related stimuli, facilitates positive reinforcement, and regulates intake of rewarding substances like alcohol and sucrose (Britt et al., 2012; Keistler et al., 2017; Millan et al., 2017; Namburi et al., 2015; Stuber et al., 2011). The BLA also projects to the bed nucleus of the stria terminalis (BNST), a brain region that regulates fear and anxiety-like behavior (Kim et al., 2013; Lange et al., 2017). Inhibiting either the BLA or the BNST using optogenetics or pharmacological inactivation reduces fear and anxiety-like behavior (Hajizadeh Moghaddam et al., 2008; Kim et al., 2013; Läck et al., 2007; Sugiyama et al., 2018; Walker & Davis, 1997; Yin et al., 2019), and yet optogenetic inhibition of the BLA projection to the BNST is anxiogenic (Kim et al., 2013). Therefore, in an effort to understand the projection-specific neurophysiological mechanisms underlying withdrawal-induced anxiety and increased alcohol intake, we compared how CIE/WD alters the neurophysiology of BLA neurons projecting to the NAC (BLA-NAC) and BNST (BLA-BNST) as representative ‘reward’ and ‘aversion’ brain regions. We found that CIE/WD enhances glutamatergic function and neuronal excitability in BLA-NAC and BLA-BNST neurons in a circuit- and sex-dependent manner. Ultimately, understanding the neurophysiological changes in these BLA projection neurons could uncover more effective pharmacotherapeutic targets aimed at reducing alcohol consumption and withdrawal-induced anxiety, as well as preventing relapse in alcohol-dependent individuals.

## EXPERIMENTAL PROCEDURES

### Animals

Male and female Sprague-Dawley rats were obtained from Envigo (Indianapolis, IN). On arrival, the rats were pair-housed and given *ad libitum* access to water and standard rat chow. Animal housing rooms are on a reverse 12-hour light-dark cycle (Lights OFF: 9 AM, Lights ON: 9 PM). Animals were approximately 5 weeks of age (130 g) when they were surgerized and approximately 11 weeks of age when electrophysiological recordings were completed. All animal care procedures are in accordance with the National Institutes of Health *Guide for the Care and Use of Laboratory Animals* and approved by the Wake Forest Animal Care and Use Committee (WF-ACUC).

### Stereotaxic surgeries

The experimental timeline is as follows (**Figure 1A**). After a brief acclimation period, the rats underwent stereotaxic microinjection surgeries as described previously (McGinnis, Parrish, & McCool, 2020). Briefly, rats were anesthetized using 3% isoflurane anesthesia and maintained under continuous 2-3% isoflurane for the remainder of the surgery with the oxygen flowmeter set at a rate of 1.0 L/min. Rats received a bilateral microinjection (0.5 µL/side) of a retrogradely transported adeno-associated viral vector expressing GFP (AAVrg-CAG-GFP; Addgene 37825-AAVrg) or tdTomato (AAVrg-CAG-tdTomato; Addgene 59642-AAVrg) into the NAC or BNST, respectively (**Figure 1B**). We used the following coordinates for the NAC and BNST injection sites (mm in relation to bregma): NAC (AP +1.80, ML ±1.50, DV 6.72) and BNST (AP -0.24, ML ±0.80, DV 6.45). A microinjection syringe pump (Harvard Apparatus) injected the viral vectors at a rate of 0.1 µL/min for 5 minutes. Injectors were left in place for 10 minutes following the microinjection to allow the virus to diffuse into the target brain region. After completing the surgery, the incision was sutured and rats received subcutaneous injections of 3 mg/kg ketoprofen for pain management and 2 mL of warmed sterile saline. The rats were single-housed until sutures were removed 6-7 days later, after which they were pair-housed for the remainder of the experiment. In total, animals were given a 4-week period to recover and allow the virus to be transported and expressed in BLA neurons prior to CIE exposure. We confirmed injection sites by collecting coronal slices containing the NAC and BNST and visualizing postmortem GFP and tdTomato expression using fluorescence microscopy (**Figure 1C**). Animals with incorrect viral placement were excluded.

**Figure 1.**
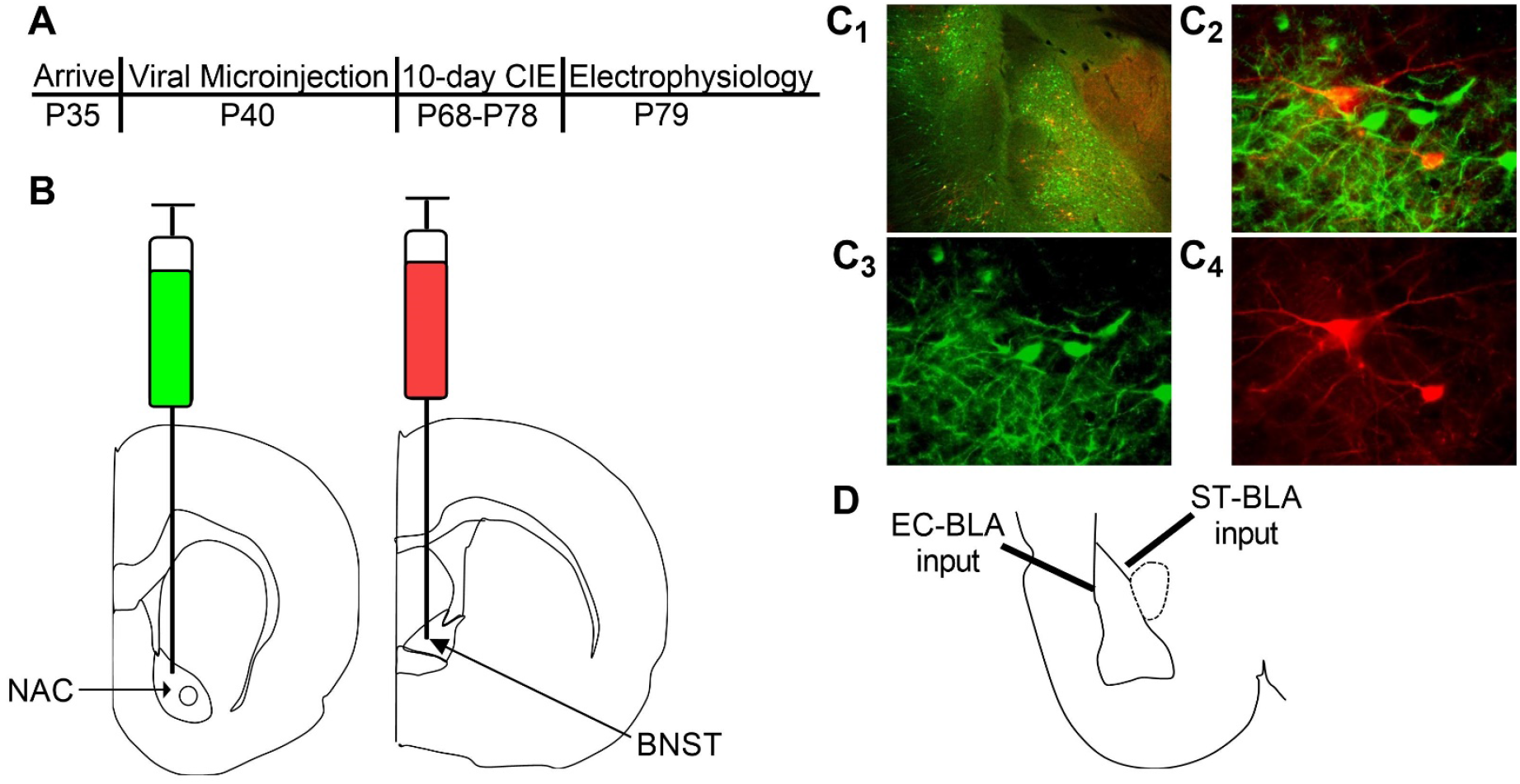
Retrograde labeling of BLA-NAC and BLA-BNST neurons. (A) Timeline of experimental procedures. (B) Schematic illustrating AAVrg-CAG-GFP into the NAC (AP +1.80 mm, ML ±1.5 mm, DV 6.72 mm) and AAVrg-CAG-tdTomato into the BNST (AP -0.24 mm, ML ±0.80 mm, DV 6.45 mm). (C_1_) 4X overlay of GFP-expressing BLA-NAC and tdTomato-expressing BLA-BNST neurons. (C_2_) 40X overlay of GFP-expressing BLA-NAC and tdTomato-expressing BLA-BNST neurons. (C_3_) GFP expression in BLA-NAC neurons under 40X magnification. (C_4_) tdTomato expression in BLA-BNST neurons under 40X magnification. (D) Schematic illustrating the placement of the stimulating electrode when activating the EC-BLA and ST-BLA inputs.

### Chronic intermittent ethanol and withdrawal (CIE/WD)

After the 4-week recovery period, rats were exposed to CIE via vapor inhalation using our standard laboratory procedure, as described previously (McGinnis, Parrish, & McCool, 2020; McGinnis, Parrish, Chappell, et al., 2020; Morales et al., 2015, 2018). Briefly, the rats were placed in custom-built plexiglass ethanol vapor chambers (Triad Plastics, Winston-Salem, NC) with their home-cages. Ethanol vapor was produced by pushing compressed air through a bubble stone submerged in 95% ethanol. The ethanol vapor was then pumped into the vapor chamber at 20-25 mg/dL. Rats were exposed to ethanol vapor for 12 hours per day for 10 consecutive days followed by a final 24-hour withdrawal period. A 10-day CIE exposure and 24-hr WD has been shown to model alcohol dependence in rodents by increasing anxiety-like behavior during WD and subsequently increasing alcohol intake (McGinnis, Parrish, & McCool, 2020; McGinnis, Parrish, Chappell, et al., 2020; Morales et al., 2015, 2018). Rats were weighed daily and tail blood samples were taken twice during the 10-day exposure to monitor blood ethanol concentrations (BECs; target = 150-275 mg/dL). BECs were measured using a standard, commercially-available alcohol dehydrogenase/NADH (nicotinamide adenine dinucleotide plus hydrogen) enzymatic assay (Carolina Liquid Chemistries). Average BECs during the CIE exposure were 224.3 ± 6.35 mg/dL in males and 221.5 ± 9.395 mg/dL in females. Air-exposed controls were housed in similar conditions except the rats only received room air.

### Electrophysiology

#### Slice preparation

Animals were anesthetized with isoflurane prior to decapitation, as stated in the animal care protocols approved by WF-ACUC. The brains were rapidly removed and allowed to incubate in an ice-cold sucrose-modified artificial cerebrospinal fluid (ACSF) solution containing the following (in mM): 180 sucrose, 30 NaCl, 4.5 KCl, 1 MgCl_2_ · 6 H_2_O, 26 NaHCO_3_, 1.2 NaH_2_PO_4_, 10 _D_-glucose, and 0.10 ketamine, oxygenated with 95% O_2_ / 5% CO_2_. Coronal brain slices (250 µm) containing the BLA were prepared using a VT1200/S vibrating blade microtome (Leica, Buffalo Grove, IL) and then incubated for 1-hour prior to electrophysiology recordings in oxygenated, room-temperature, standard ACSF solution containing the following (in mM): 126 NaCl, 3 KCl, 2 MgSO_4_ · 7 H_2_O, 26 NaHCO_3_, 1.25 NaH_2_PO_4_, 10 d-glucose, and 2 CaCl_2_ · 2 H_2_O. All chemicals were obtained from Tocris (Ellisville, Missouri) or Sigma-Aldrich (St. Louis, MO).

#### Whole-cell patch-clamp recording

After the 1-hour incubation period, BLA slices were transferred to a submersion-type recording chamber that was continuously perfused with oxygenated, room-temperature, standard ACSF at a rate of 2 mL/min. Fluorescently-labeled BLA pyramidal neurons were patched using an Olympus BZ51WI infrared differential interference contrast (IR-DIC) microscope with fluorescence attachments. Voltage clamp recordings were performed using recording electrodes filled with a Cs-gluconate intracellular solution containing (in mM): 145 CsOH, 10 EGTA, 5 NaCl, 1 MgCl_2_ · 6 H_2_O, 10 HEPES, 4 Mg-ATP, 0.4 Na-GTP, 0.4 QX314, 1 CaCl_2_· 2 H2O, pH ∼7.3 (adjusted with gluconic acid), osmolarity ∼285 Osm/L (adjusted with sucrose). Current clamp recordings were performed using recording electrodes filled with a K-gluconate intracellular solution containing the following (in mM): 145 K-gluconate, 10 EGTA, 5 NaCl, 1 MgCl_2_ · 6 H_2_O, 10 HEPES, 2 Mg-ATP, 0.1 Na-GTP, pH ∼7.3 (adjusted with KOH), osmolarity ∼285 Osm/L (adjusted with H_2_O). Data were acquired using an Axopatch 700B amplifier (Molecular Devices) and analyzed using pClamp 10 software (Molecular Devices). Presumptive BLA pyramidal neurons were included based on visual and electrophysiological characteristics: high membrane capacitance (> 100 pF) and low access resistance (≤ 25 M?) for voltage-clamp recordings, and resting membrane potential (≤ -55 mV) and action potentials that overshoot 0 mV for current-clamp recordings. Cells that did not meet these criteria were excluded from data analysis.

#### Voltage-clamp recordings

Glutamatergic excitatory postsynaptic currents (EPSCs), were recorded at a holding membrane potential of -65 mV and pharmacologically isolated using picrotoxin (100 µM; GABA_A_ receptor antagonist) in the bath ACSF. Glutamatergic synaptic responses were electrically evoked every 30 seconds by placing a concentric bipolar stimulating electrode (FHC Inc, Bowdoin, ME) within the *stria terminalis* or external capsule (**Figure 1D**) as indicated below. Stimulation intensities were submaximal and normalized to elicit EPSCs with amplitudes of ∼100 pA. Presynaptic glutamatergic function was measured using paired electrical stimuli delivered to the *stria terminalis* and separated by a 50-millisecond inter-pulse interval. Using the peak amplitude of the evoked EPSCs, we calculated a paired-pulse ratio (PPR) with the following formula: (Peak 2 - Peak 1) / Peak 1. PPR with short inter-pulse intervals are inversely related to neurotransmitter release probability (Dobrunz & Stevens, 1997). Postsynaptic glutamatergic function was measured using strontium substitution. For these recordings, calcium (2 mM) in the bath ACSF was replaced with strontium (2 mM) and a single electrical stimulus was delivered to the external capsule. Strontium is a poor substitute for calcium in terms of calcium-dependent neurotransmitter release and thus a single electrical stimulus will elicit an initial burst of neurotransmitter release followed by prolonged asynchronous neurotransmitter release (S. Choi & Lovinger, 1997). We measured the amplitude and inter-event interval of the asynchronous EPSCs (aEPSCs) occurring in the 400-ms window beginning 50 ms after the stimulus using MiniAnalysis software and then calculated the median amplitude and inter-event interval for each neuron. The amplitude of aEPSCs is related to AMPA receptor function and therefore postsynaptic glutamatergic function, whereas the inter-event interval of aEPSCs has an inverse relationship with calcium-independent presynaptic glutamatergic function (S. Choi & Lovinger, 1997).

#### Current-clamp recordings

Neuronal excitability and passive properties were examined in the whole-cell configuration using 600-msec current injections ranging from -100 pA to +300 pA. Current step injections increased in 25 pA increments; and steps were separated by 20 second inter-trial intervals. This experiment measured passive membrane properties (resting membrane potential and input resistance), evoked action potentials, and the fast and medium afterhyperpolarization (fAHP and mAHP). Resting membrane potential was assessed prior to the first current step and input resistance was calculated as the ratio of peak voltage deflection to the amount of current injected using current steps that did not elicit an action potential. Excitability measurements were analyzed by plotting the number of action potentials elicited at each current step. The fAHP and mAHP amplitudes were measured using the first action potential elicited on the lowest current step for each neuron. The fAHP and mAHP amplitudes were measured as previously described (Rau et al., 2015). We also used a 600-msec current ramp beginning at -100 pA and ending at +300 pA. This protocol was repeated for a total of 5 trials and action potential properties were derived from the first action potential elicited for each of the 5 trials. The action potential threshold was defined as the membrane potential when the action potential was elicited and the rheobase current was the corresponding current injected as the membrane potential reached the threshold. We also measured the amplitude, half-width, rise time, rise slope (10%-90%), decay time, and decay slope (90%-10%) of the first action potential (pClamp 10 software, Molecular Devices). These two recording paradigms were conducted under two conditions. In the first set of experiments, we only added picrotoxin (100 µM) to the bath ACSF. In the second set, we measured intrinsic excitability by blocking all synaptic transmission with picrotoxin (100 µM), APV (50 µM; NMDA receptor antagonist), and DNQX (20 µM; AMPA/kainate receptor antagonist) added to the bath ACSF.

### Statistical Analysis

Statistical analyses were conducted using Prism 9 (GraphPad Software). Data from BLA-NAC and BLA-BNST neurons were analyzed separately with 2-way ANOVAs or mixed-effects analyses, as indicated. Main effects and interactions were examined further with Bonferroni *post hoc* analyses to determine the locus of the effect. A value of p<0.05 was considered to be statistically significant. All data are presented as mean ± SEM.

## RESULTS

### BLA-NAC and BLA-BNST neurons can be identified with retrograde labeling and fluorescence microscopy

AAVrg viral particles expressing GFP or tdTomato were injected into the NAC or BNST, respectively (**Figure 1B**). Approximately 6 weeks after the microinjections, we examined postmortem GFP and tdTomato expression in BLA neurons (**Figure 1C**). The AAVrg viral vectors produced robust expression of GFP in BLA-NAC neurons and tdTomato expression in BLA-BNST neurons. BLA-NAC and BLA-BNST neurons were located primarily in the basal nucleus of the BLA and thus all of the electrophysiological recordings reflect changes in those neurons.

### Withdrawal increases glutamate release from *stria terminalis* onto BLA-BNST neurons

After confirming that we could use retrograde labeling to identify BLA-NAC and BLA-BNST neurons, we began to characterize the effects of CIE/WD on each population of neurons. First, we examined how CIE/WD affects glutamate release probability from *stria terminalis* inputs. Numerous studies have illustrated that CIE/WD increases glutamate release from *stria terminalis* inputs onto BLA neurons overall by measuring changes in PPR (Christian et al., 2013; McGinnis, Parrish, Chappell, et al., 2020; Morales et al., 2018; Sizer et al., 2021). PPR is inversely related to neurotransmitter release probability such that a decrease in PPR indicates an increase in neurotransmitter release probability (Dobrunz & Stevens, 1997). The current study investigated whether presynaptic changes at *stria terminalis* inputs occurred in both BLA-NAC and BLA-BNST neurons. A 2-way ANOVA found no main effects of CIE/WD or sex and no interaction effect on the PPR in BLA-NAC neurons (**Figure 2A,C**). However, in BLA-BNST neurons, a 2-way ANOVA revealed a significant main effect of CIE/WD (**Figure 2B,D**; F(1,35)=18.74, p=0.0001) on the PPR with no main effect of sex or an interaction effect. The CIE/WD-induced decrease in PPR suggests that CIE/WD increases glutamate release from *stria terminalis* inputs specifically onto BLA-BNST neurons, regardless of sex.

**Figure 2.**
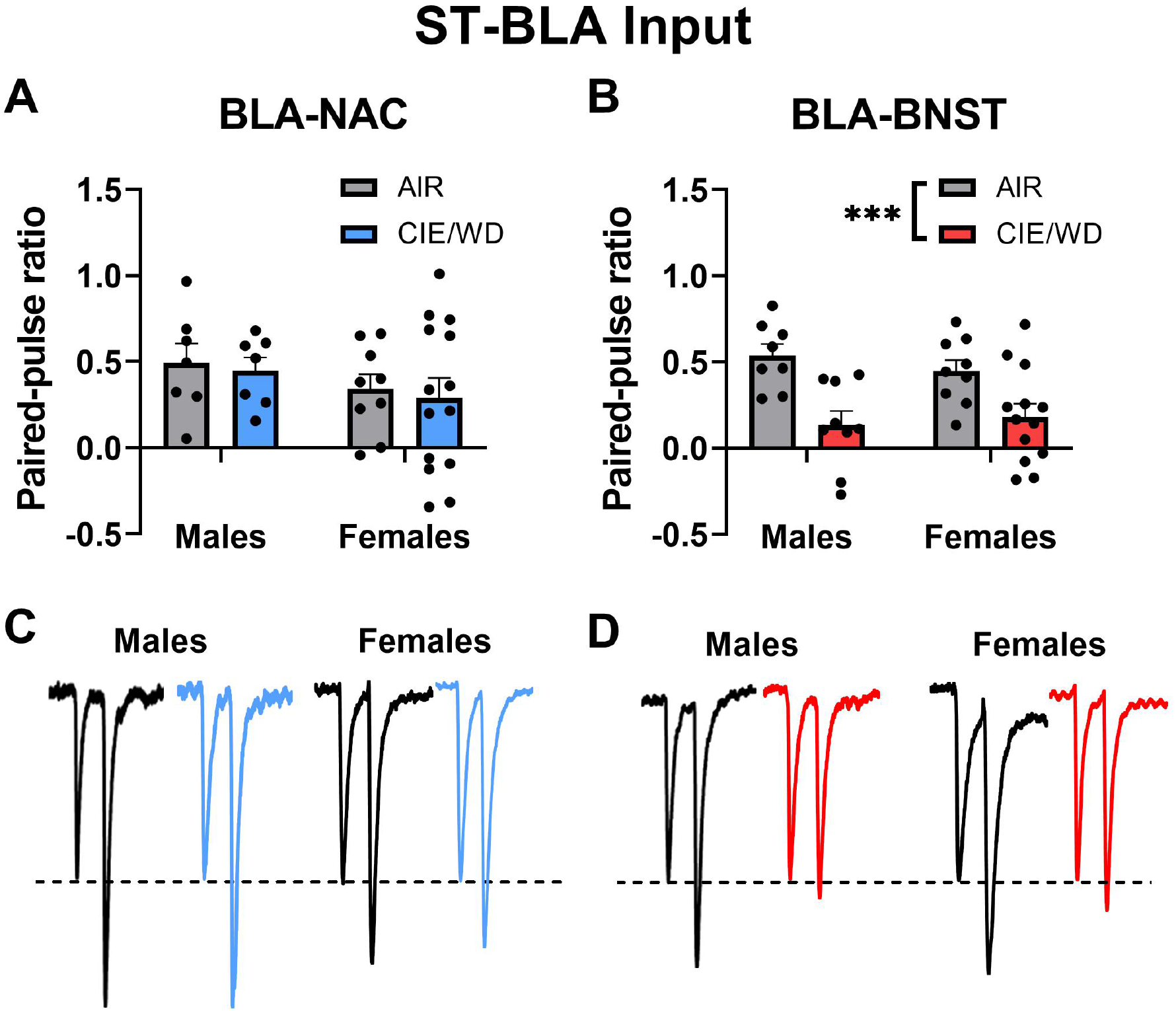
Increased glutamate release from *stria terminalis* inputs onto BLA-BNST neurons. (A) CIE/WD did not affect the PPR in BLA-NAC neurons. (B) CIE/WD decreased the PPR in BLA-BNST neurons, suggesting an increase in glutamate release from *stria terminalis* inputs onto BLA-BNST neurons. (C) Exemplar traces from paired electrical stimuli in BLA-NAC neurons from AIR males, CIE/WD males, AIR females, and CIE/WD females (left to right). (D) Exemplar traces from paired electrical stimuli in BLA-BNST neurons in AIR males, CIE/WD males, AIR females, and CIE/WD females (left to right). Data analyzed by 2-way ANOVAs. ***p<0.001

### Withdrawal increases postsynaptic glutamatergic function in external capsule synapses in a circuit- and sex-dependent manner

Prior studies have demonstrated that CIE/WD increases postsynaptic glutamatergic function at external capsule synapses with BLA neurons overall (Christian et al., 2012; McGinnis, Parrish, & McCool, 2020; Morales et al., 2018). We examined how CIE/WD affects postsynaptic function at external capsule synapses with different projection-specific populations of BLA neurons by measuring the amplitude of aEPSCs (Choi & Lovinger, 1997). In BLA-NAC neurons, a 2-way ANOVA found significant main effects of CIE/WD (**Figure 3A,E**; F(1,33)=7.073, p=0.0120) and sex (F(1,33)=4.162, p=0.0494) on aEPSC amplitude but no interaction effect. A post-hoc test revealed that CIE/WD only increased aEPSC amplitude in male BLA-NAC neurons (p=0.0031), suggesting that CIE/WD increases postsynaptic glutamatergic function in external capsule synapses with male BLA-NAC neurons. In BLA-BNST neurons, a 2-way ANOVA revealed a significant main effect of CIE/WD (**Figure 3B,F**; F(1,28)=11.75, p=0.0019) and a significant CIE/WD x sex interaction effect (F(1,28)=5.288, p=0.0291), but no main effect of sex. Post-hoc tests indicated that CIE/WD specifically increased aEPSC amplitude in females (p=0.0013) and thus females are more susceptible to the postsynaptic glutamatergic changes at these synapses compared to males.

**Figure 3.**
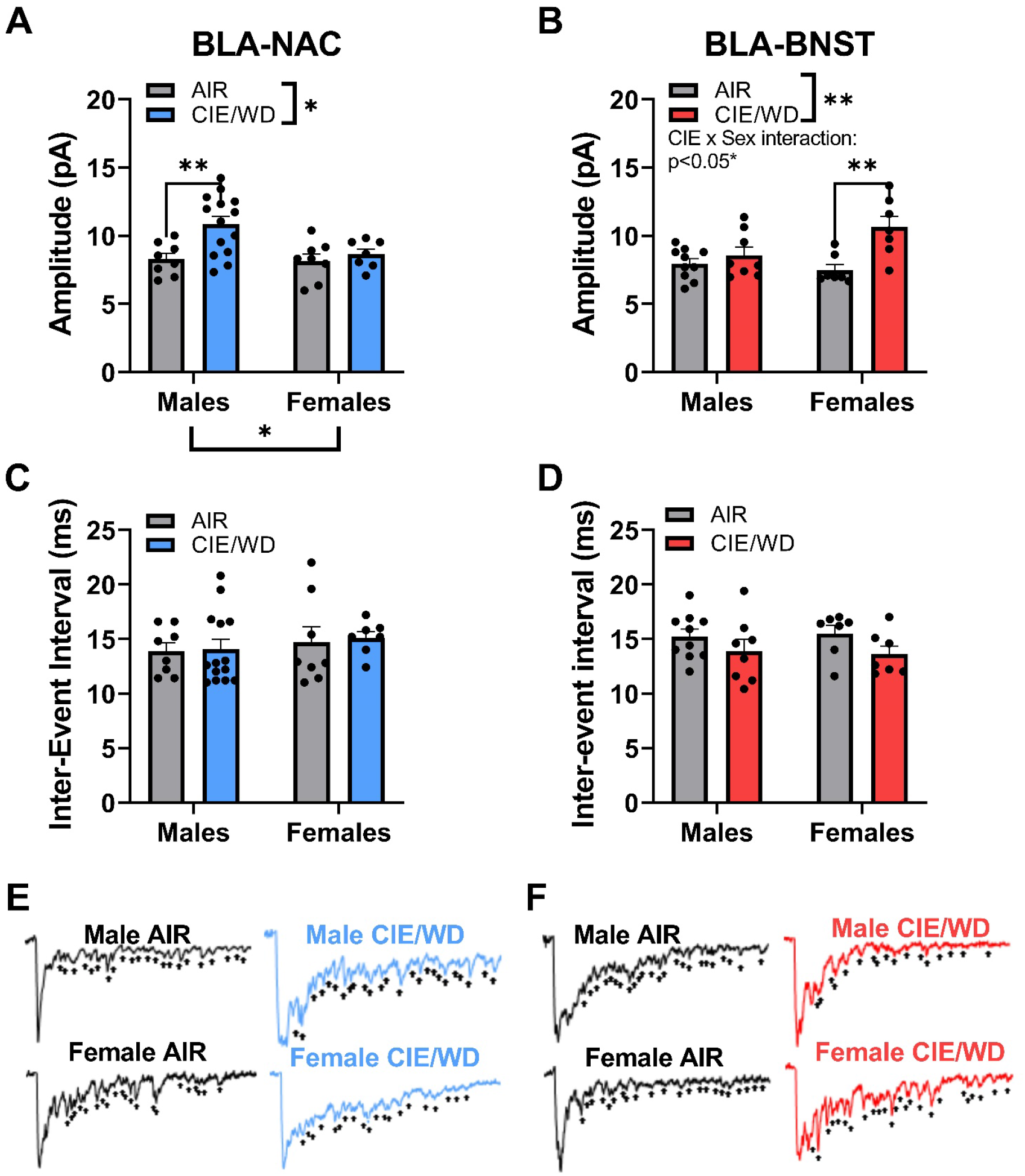
Increased glutamatergic function in external capsules synapses. (A) CIE/WD increased aEPSC amplitude in male BLA-NAC neurons. (B) CIE/WD increased aEPSC amplitude in female BLA-BNST neurons. (C) No effect of CIE/WD on aEPSC inter-event interval in BLA-NAC neurons. (D) No effect of CIE/WD on aEPSC inter-event interval in BLA-BNST neurons. (E) Exemplar traces of aEPSCs from BLA-NAC neurons in AIR males, CIE/WD males, AIR females, and CIE/WD females (+ indicate aEPSC events). (F) Exemplar traces of aEPSCs from BLA-BNST neurons in AIR males, CIE/WD males, AIR females, and CIE/WD females (+ indicate aEPSC events). Data analyzed by 2-way ANOVAs. **p<0.01, *p<0.05

The inter-event interval of aEPSCs is inversely related to calcium-independent presynaptic function (S. Choi & Lovinger, 1997). Previously, neither males nor females have shown presynaptic glutamatergic changes at external capsule synapses after CIE/WD (Christian et al., 2012; McGinnis, Parrish, & McCool, 2020; Morales et al., 2018). As expected, a 2-way ANOVA found no significant main effects and no interaction effect on the inter-event interval in BLA-NAC neurons (**Figure 3C,E**). Likewise, there were no significant main effects or an interaction effect on the inter-event interval in BLA-BNST neurons (**Figure 3D,F**). These findings confirm previous findings that CIE/WD does not alter calcium-independent presynaptic glutamatergic function at external capsule synapses (Christian et al., 2012; McGinnis, Parrish, & McCool, 2020), including BLA-NAC or BLA-BNST neurons.

### Withdrawal does not alter intrinsic excitability in BLA-NAC and BLA-BNST neurons

Chronic ethanol alters the excitability of neurons in a variety of brain regions, including the NAC and BNST (Hopf et al., 2010, 2011; Kash et al., 2009; Marty & Spigelman, 2012; Padula et al., 2015; Pati et al., 2020; Pleil et al., 2015, 2016; Shan et al., 2019), but similar experiments have not been conducted in the BLA. To determine whether CIE/WD alters the intrinsic excitability of BLA-NAC and BLA-BNST neurons, we first measured excitability with both glutamate and GABA transmission blocked. In BLA-NAC neurons, a 2-way ANOVA showed a main effect of current (**Figure 4A,C**; F(16,336)=116.0, p<0.0001) but no main effect of CIE/WD or an interaction effect. Similarly, there was a main effect of current (**Figure 4B,D**; F(1.431,27.18)=81.67, p<0.0001) but no main effect of CIE/WD or an interaction effect in BLA-BNST neurons. Thus, CIE/WD had no effect on the intrinsic excitability of BLA-NAC and BLA-BNST neurons in the absence of all synaptic transmission.

**Figure 4.**
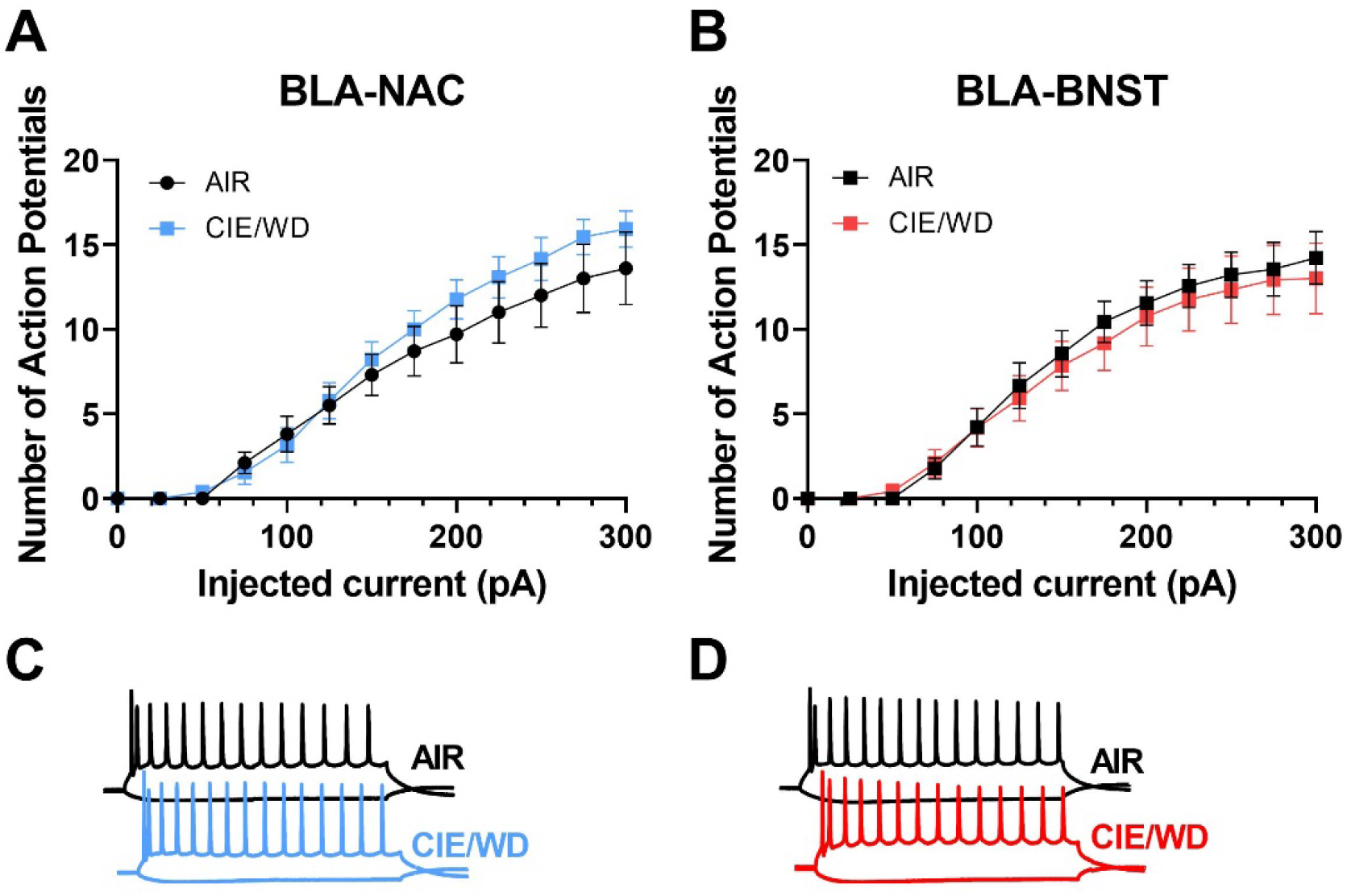
Intrinsic excitability in BLA neurons during withdrawal in males. (A) CIE/WD did not affect intrinsic excitability in BLA-NAC neurons. (B) CIE/WD did not affect intrinsic excitability in BLA-BNST neurons. (C) Exemplar traces of 600-msec current steps of -100 pA and +300 pA in BLA-NAC AIR (top) and CIE/WD (bottom) neurons. (D) Exemplar traces of 600-msec current steps of -100 pA and +300 pA in BLA-BNST AIR (top) and CIE/WD (bottom) neurons. Data analyzed by two-way ANOVAs.

### Withdrawal increases excitability in BLA-NAC and BLA-BNST neurons through distinct mechanisms

Since CIE/WD enhanced glutamatergic function in both BLA-NAC and BLA-BNST neurons, we investigated how the increase in glutamatergic function alters the excitability of these neurons when glutamatergic transmission was intact but GABAergic transmission was blocked with the GABA_A_ receptor antagonist picrotoxin. Using mixed-effects analysis with CIE/WD, sex, and current as main factors, we found significant main effects of sex (**Figure 5A,B**; F(1,49)=22.41, p<0.0001) and current (F(1.714,80.98)=245.0, p<0.0001) in BLA-NAC neurons, as well as a sex x current interaction effect (F(16,756)=16.49, p<0.0001). Although we did not find a main effect of CIE/WD, there was a significant CIE/WD x sex x current interaction effect (F(16,756)=2.526, p=0.0009) and a trending CIE/WD x sex interaction effect (F(1,49)=3.804, p=0.0569). To determine the locus of the effect, we performed mixed-effects analyses on males and females separately. In each analysis, CIE/WD and current were the main factors. In males, the mixed-effects analysis found a significant main effect of current (F(1.311,29.18)=45.90, p<0.0001) and a CIE/WD x current interaction effect (F(16,356)=1.927, p=0.0172) such that CIE/WD-exposed male rats exhibited higher BLA-NAC excitability than air-exposed controls. In contrast, there was a main effect of current (F(2.072,51.79)=308.1, p<0.0001) in female rats, but no main effect of CIE/WD or an interaction effect, indicating CIE/WD did not alter BLA-NAC excitability in females. Altogether, this suggests that BLA-NAC neuron excitability is higher in females compared to males and that CIE/WD specifically increases BLA-NAC neuron excitability in males. Finally, we used mixed-effects analysis with CIE/WD, sex, and current as main factors to examine excitability in BLA-BNST neurons (**Figure 5C,D**). We found significant main effects of CIE/WD (F(1,47)=5.014, p=0.0299) and current (F(1.732,81.31)=374.6, p<0.0001), as well as a significant CIE/WD x current interaction effect (F(16,751)=3.986, p<0.0001). These data indicate that CIE/WD increases the excitability of BLA-BNST neurons in both males and females.

**Figure 5.**
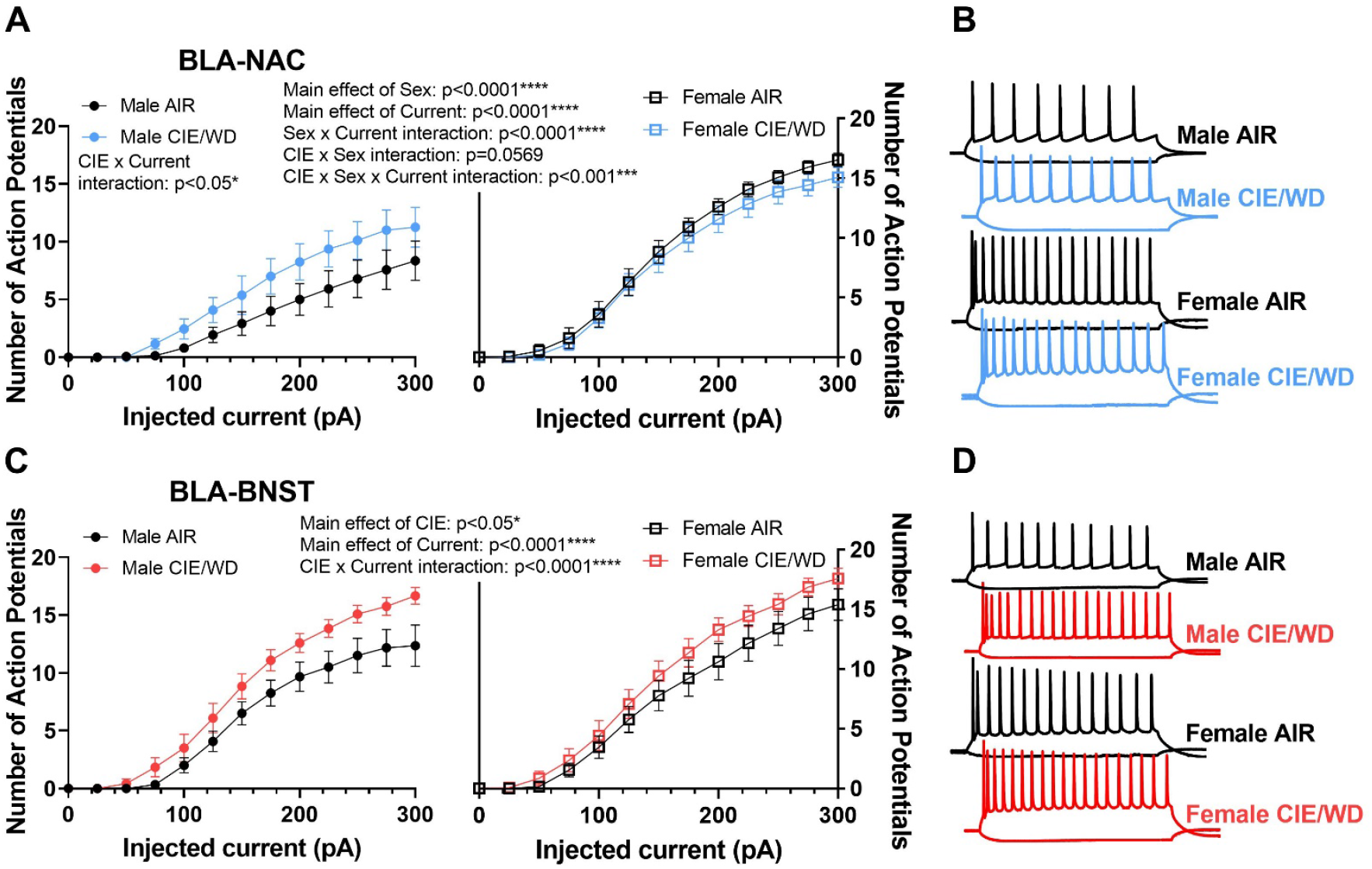
Enhanced excitability in BLA neurons during withdrawal. (A) CIE/WD increased excitability in male BLA-NAC neurons (p<0.05; left) and females had higher BLA-NAC excitability than males (p<0.0001; right). (B) Exemplar traces of 600-msec current steps of -100 pA and +300 pA in BLA-NAC neurons from AIR males, CIE/WD males, AIR females, and CIE/WD females (top to bottom). (C) CIE/WD increased excitability in BLA-BNST neurons, regardless of sex (p<0.0001). (D) Exemplar traces of 600-msec current steps of -100 pA and +300 pA in BLA-BNST neurons from AIR males, CIE/WD males, AIR females, and CIE/WD females (top to bottom). Data analyzed by mixed-effects analyses.

To determine the mechanisms driving enhanced excitability of BLA-NAC neurons in males and BLA-BNST neurons in both sexes, we first measured the input resistance, rheobase current, and action potential threshold. 2-way ANOVAs revealed a main effect of CIE/WD on input resistance in BLA-NAC neurons (**Figure 6A**; F(1,49)=7.889, p=0.0071) and BLA-BNST neurons (**Figure 6B**; F(1,46)=13.41, p=0.0006) such that input resistance increased in both BLA-NAC and BLA-BNST neurons. In contrast, CIE/WD effects on the rheobase current were projection-and sex-specific. 2-way ANOVAs in BLA-NAC neurons found significant main effects of CIE/WD (**Figure 6C**; F(1,42)=5.309, p=0.0262) and sex (F(1,42)=16.82, p=0.0002) on the rheobase current, as well as a CIE/WD x sex interaction effect (F(1,42)=8.270, p=0.0063). Post-hoc analyses revealed that CIE/WD specifically decreases the rheobase current in males (p=0.0032). These data indicate that females have a lower rheobase current than males in BLA-NAC neurons and only males exhibit a decrease in rheobase following CIE/WD. There were no significant effects on the rheobase current in BLA-BNST neurons (**Figure 6D**). Interestingly, there is a significant main effect of sex on action potential threshold in both BLA-NAC (**Figure 6E**; F(1,42)=10.28, p=0.0026) and BLA-BNST (**Figure 6F**; F(1,39)=14.81, p=0.0004) neurons such that females have a lower action potential threshold than males, but no main effect of CIE/WD or an interaction effect.

**Figure 6.**
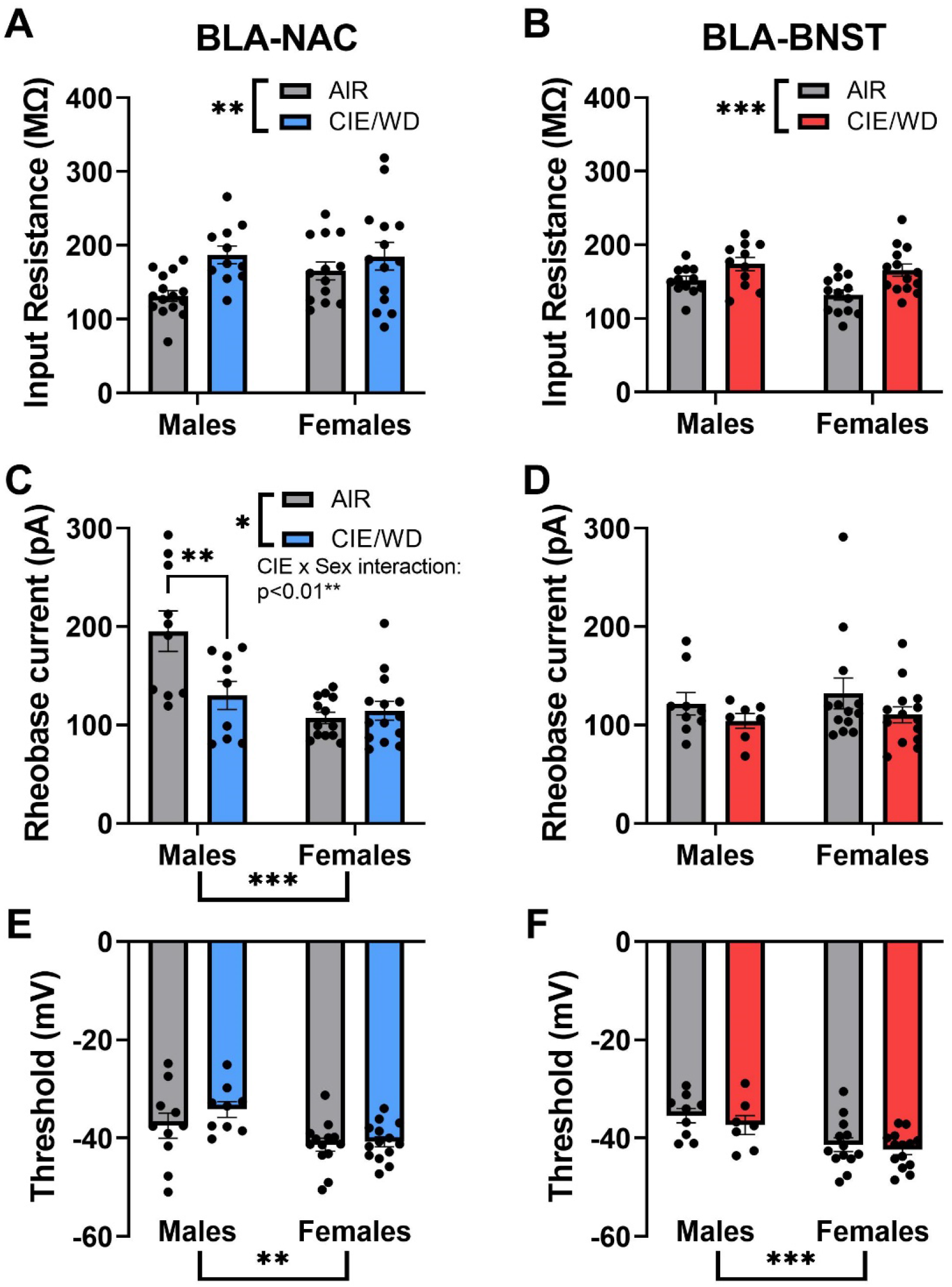
Circuit-dependent mechanisms of enhanced excitability. (A) CIE/WD increased input resistance in BLA-NAC neurons. (B) CIE/WD increased input resistance in BLA-BNST neurons. (C) Female BLA-NAC neurons had a lower rheobase current than males and CIE/WD decreased the rheobase current in male BLA-NAC neurons. (D) CIE/WD did not affect the rheobase current in BLA-BNST neurons. (E) Female BLA-NAC neurons had a lower action potential threshold than males. (F) Female BLA-BNST neurons had a lower action potential threshold than males. Data analyzed by 2-way ANOVAs. ***p<0.001, **p<0.01, *p<0.05

Both BLA-NAC and BLA-BNST neurons exhibit increased input resistance, indicating that CIE/WD closes ion channels. Increased input resistance is associated with hyperexcitability in these neurons, suggesting that the hyperexcitability is driven by a loss of hyperpolarizing current. The closing of calcium-activated potassium channels, including big conductance BK channels and small conductance SK channels, would have a similar effect. Therefore, we measured fAHP and mAHP amplitudes to determine if the CIE/WD-mediated hyperexcitability could be facilitated by BK and SK channels, respectively. CIE/WD did not impact the fAHP in BLA-NAC neurons (**Figure 7A**), but in BLA-BNST neurons, there was a significant CIE/WD x sex interaction effect (**Figure 7B**; F(1,45)=5.974, p=0.0185) and a trending CIE/WD effect (F(1,45)=3.604, p=0.0640). Post-hoc analyses illustrated that CIE/WD decreases the fAHP amplitude in male BLA-BNST neurons (p=0.0093). CIE/WD did not alter the mAHP in either BLA-NAC or BLA-BNST neurons, but there is a significant main effect of sex on the mAHP in BLA-NAC neurons (**Figure 7C**; F(1,49)=6.736, p=0.0124) and a trending effect of sex in BLA-BNST neurons (**Figure 7D**; F(1,45)=3.476, p=0.0688). This suggests that females have a larger mAHP amplitude compared to males. There were no significant effects of CIE/WD or sex on the resting membrane potential or action potential characteristics including amplitude, half-width, rise time and slope, and decay time and slope in either group of neurons (**Table 1**).

**Figure 7.**
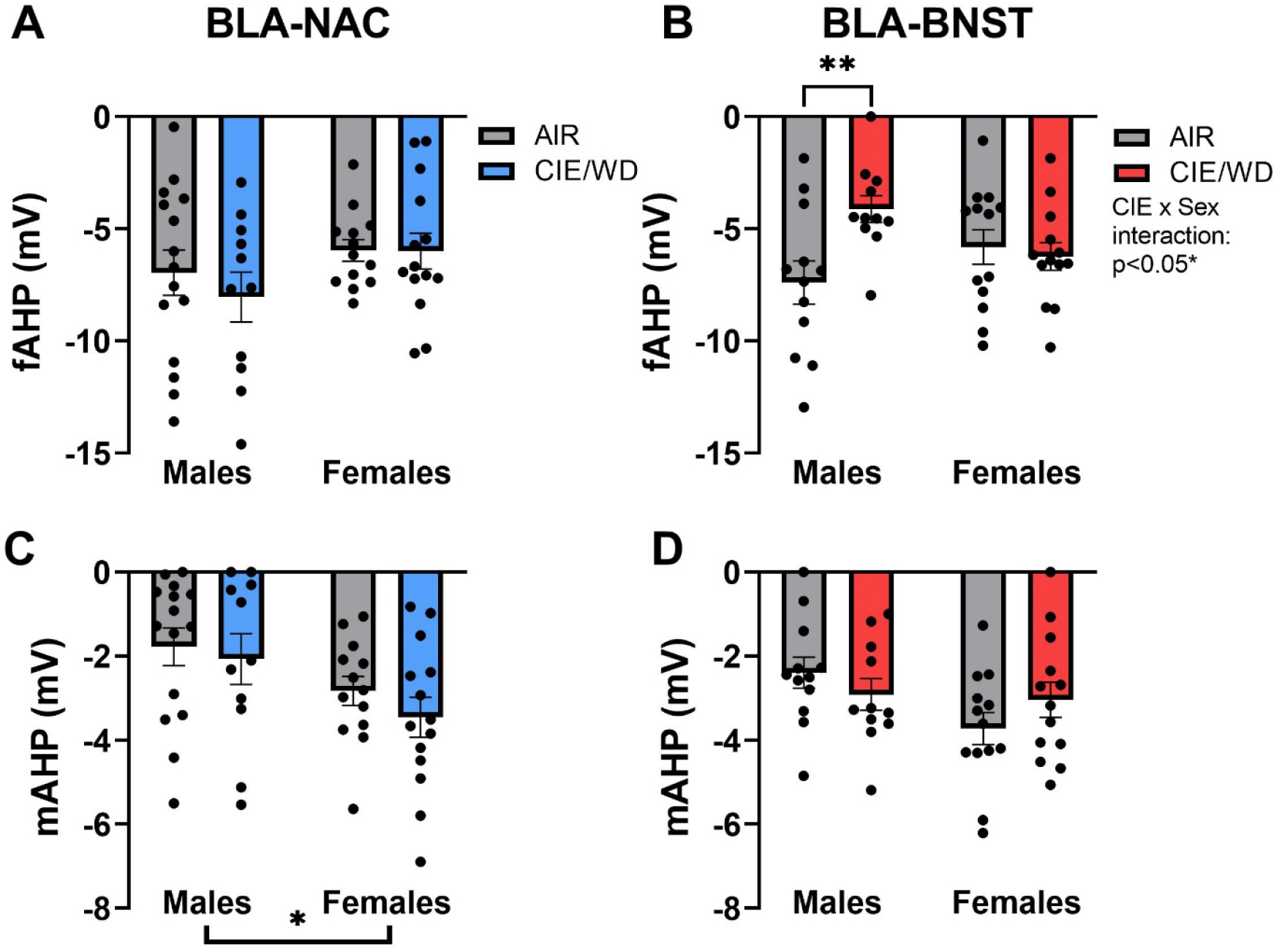
Ethanol-induced and sex-dependent effects on afterhyperpolarization in BLA neurons. (A) CIE/WD did not alter the fAHP amplitude in BLA-NAC neurons. (B) CIE/WD reduced the fAHP amplitude only in male BLA-BNST neurons. (C) Female BLA-NAC neurons had a larger mAHP amplitude compared to males. (D) CIE/WD did not alter the mAHP amplitude in BLA-BNST neurons (trending sex effect p=0.0688). Data analyzed by 2-way ANOVAs. **p<0.01, *p<0.05

**Figure 8.**
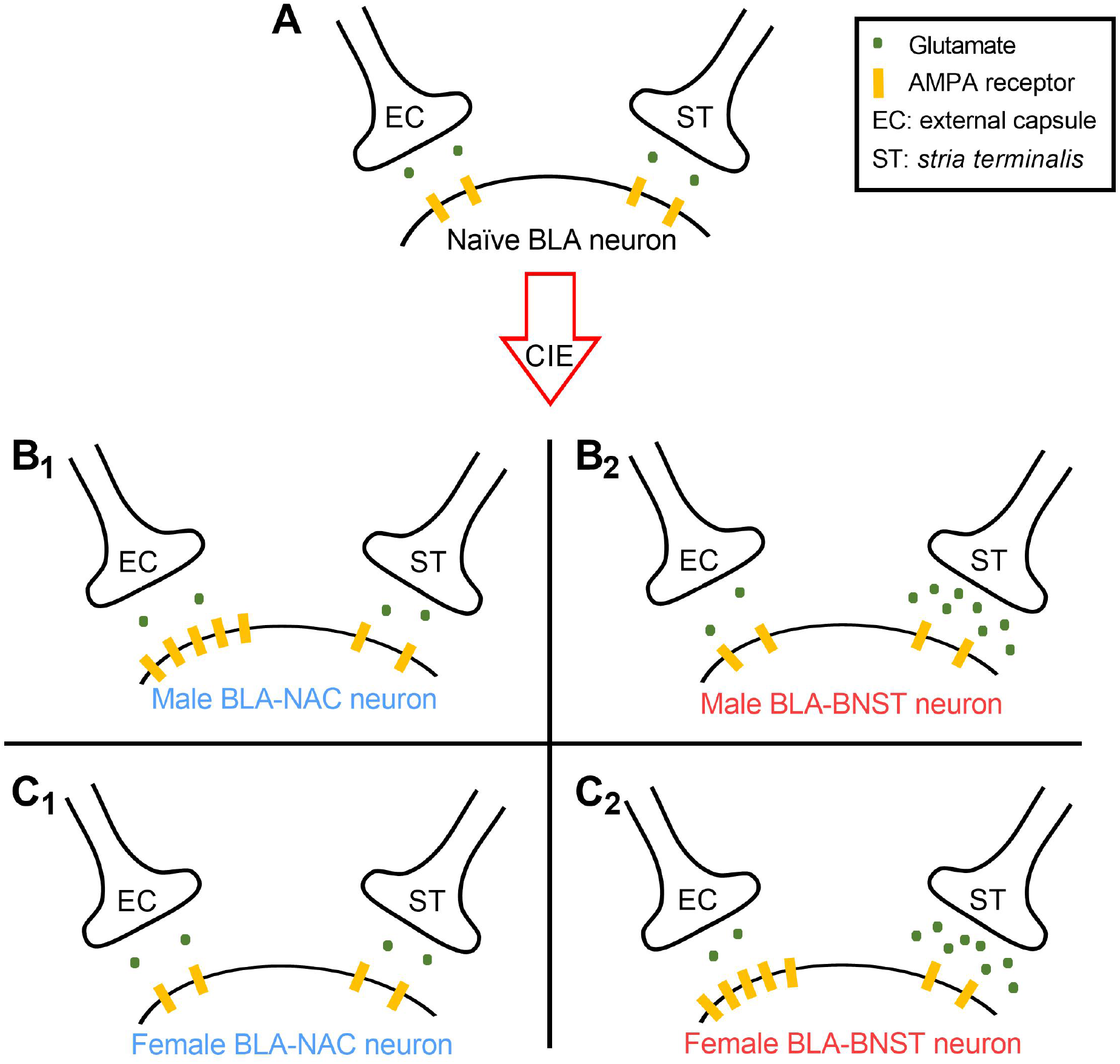
Summary findings on the effects of CIE/WD on BLA-NAC and BLA-BNST neurons at two distinct input pathways. (A) Naïve BLA synapse with EC (left) and ST (right) glutamatergic inputs. (B_1_) After CIE/WD, male BLA-NAC neurons only have increased postsynaptic function in EC synapses. (B_2_) CIE/WD increases glutamate release from ST inputs onto male BLA-BNST neurons. (C_1_) CIE/WD does not affect glutamatergic function in EC or ST synapses with female BLA-NAC neurons. (C_2_) CIE/WD increases glutamate release from ST inputs onto female BLA-BNST neurons and increases postsynaptic function in EC synapses with these neurons.

## DISCUSSION

The current study demonstrates that BLA-NAC and BLA-BNST neurons exhibit circuit- and sex-dependent neurophysiological changes following CIE/WD, suggesting that CIE/WD differentially impacts BLA projections. In BLA-NAC neurons, CIE/WD enhances postsynaptic glutamatergic function and excitability specifically in males. By comparison, the effects of CIE/WD on glutamatergic function and excitability of BLA-BNST neurons are largely independent of sex. CIE/WD increases glutamate release and excitability in both sexes, plus a female-specific increase in postsynaptic glutamatergic function.

### BLA-NAC Projection

In BLA-NAC neurons, CIE/WD increases postsynaptic glutamatergic function at external capsule synapses but has no presynaptic effects at *stria terminalis* or external capsule inputs. Interestingly, the increased postsynaptic glutamatergic function is only observed in males and facilitates a male-specific increase in BLA-NAC excitability, driven by increased input resistance and a male-specific decrease in the rheobase current. Female BLA-NAC neurons are more excitable at baseline compared to males due to a lower rheobase current and action potential threshold and do not become more excitable after CIE/WD. These data appear to parallel behavioral studies illustrating that females consume more alcohol at baseline compared to males and that only males increase alcohol consumption after CIE (Morales et al., 2015). Altogether, behavioral and electrophysiological experiments suggest that chronic ethanol only strengthens the excitability of reward circuits in males, leading to increased alcohol consumption, and that females may be protected from increased alcohol consumption due to high baseline excitability in BLA-NAC neurons.

Enhancing postsynaptic glutamatergic function and excitability in male BLA neurons projecting to the NAC could translate to greater glutamate release from BLA inputs onto NAC neurons following CIE/WD. BLA inputs have a relatively low glutamate release probability onto NAC neurons in naïve male mice (Britt et al., 2012) which may allow for a more dynamic response to CIE/WD exposure. There is considerable evidence that chronic alcohol exposure enhances the excitability of GABAergic medium spiny neurons in the NAC core of male rodents, the predominant neuron type in the NAC (Hopf et al., 2010, 2011; Shan et al., 2019). Ethanol-induced hyperexcitability in NAC core is driven by changes to calcium-activated potassium channels that suppress neuronal excitability by hyperpolarizing the cell membrane and making it more difficult to reach the action potential threshold. Chronic ethanol specifically reduces SK channel currents, SK-sensitive mAHP amplitude, and SK3 subunit protein expression in NAC core MSNs (Hopf et al., 2010; Shan et al., 2019).

CIE/WD increases excitability and input resistance in BLA-NAC neurons, indicating that the enhanced excitability is associated with fewer ion channels open on the cell membrane. This suggests that chronic ethanol reduces a hyperpolarizing current like those generated by calcium-activated potassium channels. It is unlikely that BK or SK channels are responsible for the enhanced excitability observed in male BLA-NAC neurons given that CIE/WD does not alter the fAHP or mAHP amplitude; however, another type of potassium channel is likely involved.

Our findings indicate that female BLA-NAC neurons have greater basal excitability compared to males, which seems to be ‘protective’ against the CIE/WD-mediated hyperexcitability observed in males. Although it is unclear whether NAC MSNs have greater basal excitability in females compared to males, greater basal excitability in female BLA-NAC neurons is consistent with previous reports that female BLA neurons in general have a higher basal firing rate and are more sensitive to glutamate-driven increases in firing rate compared to males (Blume et al., 2017). A higher basal firing rate and greater sensitivity to glutamate could facilitate higher basal excitability in female BLA-NAC neurons. Greater basal excitability in females could also translate to greater basal glutamate release from BLA inputs onto NAC neurons, further enhancing the excitability of the BLA-NAC circuit in naïve females and attenuating ethanol-induced hyperexcitability. To our knowledge, there are no published studies on sex differences in glutamate release from BLA inputs onto NAC neurons in naïve or ethanol-exposed animals. Interestingly, despite the lack of CIE/WD-mediated effects on SK-sensitive mAHP amplitudes, our data indicate that the mAHP amplitude is larger in female BLA-NAC neurons compared to males, suggesting there may be sex differences in SK channels. Recent studies have demonstrated sex differences in how SK channels inhibitors alter neuronal excitability after chronic stress in other brain regions like the dorsal raphe (Oliver et al., 2020). Moreover, the mAHP amplitude is sex-dependent in the entorhinal cortex layer VI, leading to sex differences in neuronal excitability (Chung & Bailey, 2019). These sex differences in SK channel function may be due to estrogen or progesterone effects. For instance, 17β-estradiol alters SK3 subunit mRNA expression in female gonadotropin-releasing hormone-expressing neurons (Bosch et al., 2013).

### BLA-BNST Projection

In BLA-BNST neurons, CIE/WD has profound effects on glutamatergic function at *stria terminalis* and external capsule synapses. CIE/WD augments glutamate release from *stria terminalis* inputs, independent of sex. In addition, females exhibit increased postsynaptic glutamatergic function at external capsule synapses with BLA-BNST neurons while males do not. The increased glutamatergic function translates to enhanced excitability in male and female BLA-BNST neurons driven by an overall increase in input resistance and a male-specific reduction in fAHP amplitude. Thus, aversion circuits are potentiated after CIE/WD, regardless of sex. These data are consistent with behavioral studies showing that chronic ethanol increases withdrawal-induced anxiety-like behavior in both sexes (Morales et al., 2015, 2018).

Although both male and female BLA-BNST neurons express enhanced excitability after CIE/WD regardless of sex, the mechanisms responsible for increasing excitability may be sex-dependent. CIE/WD increased the input resistance overall, indicating that the enhanced excitability is caused by a decrease in the number of channels that are open in both males and females. However, the male-specific reduction in fAHP amplitude suggests that CIE/WD-induced hyperexcitability could be driven by a decrease in the number of open BK channels in males but not females. BK channels are involved in producing the fAHP and evidence shows that acute restraint stress and fear conditioning similarly enhance excitability in the male lateral amygdala by reducing fAHP amplitude and BK channel expression (Guo et al., 2012; Sun et al., 2015). In contrast, the increased excitability in female BLA-BNST neurons is likely caused by decreasing the number of open potassium channels other than BK and SK channels.

Our data suggest that CIE/WD increases BLA-BNST neuron excitability, a circuit that regulates anxiety-like behavior. The BLA and BNST generally drive fear and anxiety-like behavior (Hajizadeh Moghaddam et al., 2008; Kim et al., 2013; Läck et al., 2007; Sugiyama et al., 2018; Walker & Davis, 1997; Yin et al., 2019), but optogenetic studies have shown that the BLA-BNST projection produces the opposite phenotype (Kim et al., 2013; Lange et al., 2017). The neurophysiological changes we describe could drive a compensatory mechanism to act against anxiogenic circuits activated by CIE/WD or they may reveal a shift in the function of the BLA-BNST projection. A similar mechanism dictates the function of the BLA projection to the lateral CeA. The BLA projects to two distinct subpopulations in the lateral CeA: SOM+/PKCδ- neurons and SOM-/PKCδ+ neurons. In naïve animals, the excitatory strength of BLA-CeA^SOM-^ synapses is stronger than BLA-CeA^SOM+^ synapses, facilitating anxiolysis (Cai et al., 2014; Janak & Tye, 2015; Li et al., 2013; Tye et al., 2011). Fear conditioning reverses the relative strength of these synapses by increasing glutamate release from the BLA onto CeA^SOM+^ neurons and decreasing glutamate release onto CeA^SOM-^ neurons (Li et al., 2013). Given the heterogeneity of the BNST, it’s plausible that CIE/WD causes a similar shift in synaptic strength such that the BLA-BNST projection is anxiolytic in naïve animals and anxiogenic in CIE/WD-exposed animals. Future behavioral experiments are critical to untangling the BLA-BNST circuitry responsible for regulating anxiety in naïve and CIE/WD-exposed animals.

### Motivational drivers of AUD: positive and negative reinforcement

Altogether, chronic ethanol impacts BLA-NAC neurons in males but not females and, although there are substantial effects on BLA-BNST neurons in both sexes, females have additional neurophysiological changes that the males do not. Female BLA-BNST neurons have both pre-and postsynaptic facilitation at *stria terminalis* and external capsule synapses, respectively. The disparity suggests that males are driven primarily by positive reinforcement and females are driven by negative reinforcement following CIE. Clinical studies have illustrated that adolescent boys are significantly more likely to initiate first-time alcohol consumption for alcohol enhancement motives (i.e. drinking to have fun) whereas adolescent girls are twice as likely to initiate drinking as a coping mechanism for negative emotions (Dir et al., 2017a; Kuntsche & Müller, 2011). Furthermore, binge drinking was associated with higher impulsivity/sensation-seeking in young adult men and higher neuroticism/anxiety in young adult women (Adan et al., 2016; Dir et al., 2017b). Thus, it is imperative that we consider the motivations behind risky drinking behaviors to develop pharmacotherapies that effectively target reward-seeking and negative affect for men and women, respectively.

### Implications for the timecourse of neurophysiological changes in the BLA

Previous studies examining the time course of neurophysiological changes in the BLA after CIE/WD found that presynaptic changes at *stria terminalis* synapses occur prior to postsynaptic changes at external capsule synapses, and that both pre- and postsynaptic glutamatergic changes are delayed in females compared to males (Morales et al., 2018). Given that presynaptic changes at *stria terminalis* synapses are observed in BLA-BNST neurons while postsynaptic changes at external capsule synapses are predominantly in BLA-NAC neurons, the “presynaptic prior to postsynaptic” pattern may reflect the time course in which aversion and reward circuits are affected. That would suggest that withdrawal-induced anxiety-like behavior develops prior to changes in reward-related behaviors like alcohol intake. However, in females, postsynaptic changes at external capsule synapses appear to be primarily driven by BLA-BNST neurons, suggesting that some aspects of aversion may be augmented further in females after longer durations of alcohol exposure.

### Input versus output pathway contributions

The BLA receives afferents from numerous brain regions, including but not limited to, the prefrontal cortex, temporal and insular regions, thalamus, and hippocampus (Huang et al., 2021; Leichnetz & Astruc, 1977; McDonald, 1998; McDonald et al., 1996; Sah et al., 2003). The projection targets of the BLA are equally diverse, including but not limited to, the NAC, BNST, central amygdala (CeA), prefrontal cortex, and hippocampus (Beyeler et al., 2016, 2018; Huang et al., 2021; Janak & Tye, 2015; Senn et al., 2014). A recent study utilized a novel approach to examine BLA input-output circuits, specifically the collateralization between BLA projections to distinct targets and input diversity to projection-specific BLA neuron subpopulations (Huang et al., 2021). Importantly, there is little collateralization between BLA projections to the NAC and BNST and yet, the inputs to BLA-NAC and BLA-BNST neurons are indistinguishable. These findings, in the context of our study, highlight the specificity of CIE/WD effects on BLA-NAC and BLA-BNST neurons. Although CIE/WD ultimately enhanced excitability in both groups of neurons, the mechanisms by which this occurred are distinct despite receiving identical inputs that each have input-specific effects from CIE/WD. BLA-NAC neurons only experienced postsynaptic facilitation, whereas BLA-BNST neurons exhibited presynaptic facilitation in both sexes, as well as postsynaptic facilitation in females.

## Conclusions

These findings are the first to demonstrate projection-specific alterations in BLA neurophysiology following CIE/WD. BLA-mediated behaviors, including anxiety-like behavior and alcohol consumption, are initiated through downstream projections to anxiety and reward-related regions like the BNST and NAC, respectively. Therefore, understanding how CIE/WD changes glutamatergic function and the excitability of projection-specific populations of BLA neurons is imperative to identifying the neurophysiological mechanisms driving WD-induced anxiety and increased alcohol intake. BLA-NAC (‘reward’) neurons exhibit postsynaptic facilitation of glutamatergic function and hyperexcitability exclusively in CIE-exposed males. Moreover, female BLA-NAC neurons have higher basal excitability compared to males. The striking parallels between the baseline sex differences and CIE-induced effects on BLA-NAC excitability and alcohol intake strongly suggest that neurophysiological changes in BLA-NAC neurons drive alcohol intake. CIE/WD effects in BLA-BNST (‘aversion’) neurons are largely independent of sex as they exhibit presynaptic facilitation of glutamatergic function and hyperexcitability in both sexes, as well as postsynaptic facilitation of glutamatergic function in females. The parallels between sex-independent, CIE-induced increases in BLA-BNST excitability and anxiety-like behavior strongly suggest that neurophysiological changes in BLA-BNST neurons drive anxiety-like behavior. Further experiments will be necessary to fully elucidate the mechanisms by which females are protected from CIE/WD-induced changes to glutamatergic function and excitability. In addition, future experiments could explore other BLA projections that regulate anxiety and reward-related behaviors, such as projections to the CeA, hippocampus, and prelimbic cortex.

## Acknowledgements

This work is supported by NIH/NIAAA grants T32 AA007565 (MEP), P50 AA026117 (BAM), and R01 AA014445 (BAM).

## REFERENCES

Adan, A., Navarro, J. F., & Forero, D. A. (2016). Personality profile of binge drinking in university students is modulated by sex. A study using the Alternative Five Factor Model. Drug and Alcohol Dependence, 165, 120–125. https://doi.org/10.1016/j.drugalcdep.2016.05.015

Almeida, O. F., Shoaib, M., Deicke, J., Fischer, D., Darwish, M. H., & Patchev, V. K. (1998). Gender differences in ethanol preference and ingestion in rats. The role of the gonadal steroid environment. The Journal of Clinical Investigation, 101(12), 2677–2685. https://doi.org/10.1172/JCI1198

Amodeo, L. R., Wills, D. N., Sanchez-Alavez, M., Nguyen, W., Conti, B., & Ehlers, C. L. (2018). Intermittent voluntary ethanol consumption combined with ethanol vapor exposure during adolescence increases drinking and alters other behaviors in adulthood in female and male rats. Alcohol, 73, 57–66. https://doi.org/10.1016/j.alcohol.2018.04.003.ALCOHOL

Beyeler, A., Chang, C.-J., Silvestre, M., Levegue, C., Namburi, P., Wildes, C. P., & Tye, K. M. (2018). Organization of valence-encoding and projection-defined neurons in the basolateral amygdala. Cell Rep, 22(4), 905–918. https://doi.org/10.1016/j.physbeh.2017.03.040

Beyeler, A., Namburi, P., Glober, G. F., Simonnet, C., Gwendolyn, G., Conyers, G. F., Luck, R., Wildes, C. P., & Tye, K. M. (2016). Divergent routing of positive and negative information from the amygdala during memory retrieval. Neuron, 90(2), 348–361. https://doi.org/10.1016/j.neuron.2016.03.004.Divergent

Blume, S. R., Freedberg, M., Vantrease, J. E., Chan, R., Padival, M., Record, M. J., De Joseph, M. R., Urban, J. H., & Rosenkranz, J. A. (2017). Sex-And estrus-dependent differences in rat basolateral amygdala. Journal of Neuroscience, 37(44), 10567–10586. https://doi.org/10.1523/JNEUROSCI.0758-17.2017

Bosch, M. A., Tonsfeldt, K. J., & Rønnekleiv, O. K. (2013). mRNA expression of ion channels in GnRH neurons: Subtype-specific regulation by 17β-estradiol. In Molecular and Cellular Endocrinology (Vol. 367, Issues 1–2). https://doi.org/10.1016/j.mce.2012.12.021.mRNA

Britt, J. P., Benaliouad, F., McDevitt, R. A., Stuber, G. D., Wise, R. A., & Bonci, A. (2012). Synaptic and behavioral profile of multiple glutamatergic inputs to the Nucleus Accumbens. Neuron, 76(4), 790–803. https://doi.org/10.1016/j.neuron.2012.09.040.Synaptic

Cai, H., Haubensak, W., Anthony, T., & Anderson, D. J. (2014). Central amygdala PKC-δ+ neurons mediate the influence of multiple anorexigenic signals. Nature Neuroscience, 17(9), 1240–1248. https://doi.org/10.1038/nn.3767.Central

Choi, J. M., Padmala, S., Spechler, P., & Pessoa, L. (2013). Pervasive competition between threat and reward in the brain. Social Cognitive and Affective Neuroscience, 9(6), 737–750. https://doi.org/10.1093/scan/nst053

Choi, S., & Lovinger, D. M. (1997). Decreased frequency but not amplitude of quantal synaptic responses associated with expression of corticostriatal long-term depression. Journal of Neuroscience, 17(21), 8613–8620. https://doi.org/10.1523/jneurosci.17-21-08613.1997

Christian, D. T., Alexander, N. J., Diaz, M. R., & McCool, B. A. (2013). Thalamic glutamatergic afferents into the rat basolateral amygdala exhibit increased presynaptic glutamate function Following Withdrawal from Chronic Intermittent Ethanol. Neuropharmacology, 65, 134–142. https://doi.org/10.1016/j.neuropharm.2012.09.004.Thalamic

Christian, D. T., Alexander, N. J., Diaz, M. R., Robinson, S., & McCool, B. A. (2012). Chronic intermittent ethanol and withdrawal differentially modulate basolateral amygdala AMPA-type glutamate receptor function and trafficking. Neuropharmacology, 62(7), 2429–2438. https://doi.org/10.1016/j.neuropharm.2012.02.017

Chung, B. Y. T., & Bailey, C. D. C. (2019). Sex differences in the nicotinic excitation of principal neurons within the developing hippocampal formation. Developmental Neurobiology, 79(2), 110–130. https://doi.org/10.1002/dneu.22646

Diaz, M. R., Chappell, A. M., Christian, D. T., Anderson, N. J., & McCool, B. A. (2011). Dopamine D3-like receptors modulate anxiety-like behavior and regulate gabaergic transmission in the rat lateral/basolateral amygdala. Neuropsychopharmacology, 36, 1090–1103. https://doi.org/10.1038/npp.2010.246

Dir, A. L., Bell, R. L., Adams, Z. W., & Hulvershorn, L. A. (2017a). Gender Differences in Risk Factors for Adolescent Binge Drinking and Implications for Intervention and Prevention. Frontiers in Psychiatry, 8(December). https://doi.org/10.3389/fpsyt.2017.00289

Dir, A. L., Bell, R. L., Adams, Z. W., & Hulvershorn, L. A. (2017b). Gender Differences in Risk Factors for Adolescent Binge Drinking and Implications for Intervention and Prevention. Frontiers in Psychiatry, 8(December). https://doi.org/10.3389/fpsyt.2017.00289

Dobrunz, L. E., & Stevens, C. F. (1997). Heterogeneity of release probability, facilitation, and depletion at central synapses. Neuron, 18, 995–1008. https://doi.org/10.1016/S0896-6273(00)80338-4

Guo, Y. Y., Liu, S. B., Cui, G. Bin, Ma, L., Feng, B., Xing, J. H., Yang, Q., Li, X. Q., Wu, Y. mei, Xiong, L. Z., Zhang, W., & Zhao, M. G. (2012). Acute stress induces down-regulation of large-conductance Ca 2+-activated potassium channels in the lateral amygdala. Journal of Physiology, 590(4), 875–886. https://doi.org/10.1113/jphysiol.2011.223784

Hajizadeh Moghaddam, A., Roohbakhsh, A., Rostami, P., Heidary-Davishani, A., & Zarrindast, M. R. (2008). GABA and histamine interaction in the basolateral amygdala of rats in the plus-maze test of anxiety-like behaviors. Pharmacology, 82, 59–66. https://doi.org/10.1159/000131110

Hopf, F. W., Bowers, M. S., Chang, S.-J., Chen, B. T., Martin, M., Seif, T., Cho, S. L., Tye, K., & Bonci, A. (2010). Reduced nucleus accumbens SK channel activity enhances alcohol seeking during abstinence. Neuron, 65(5), 682–694. https://doi.org/10.1016/j.neuron.2010.02.015.Reduced

Hopf, F. W., Simms, J. A., Chang, S.-J., Seif, T., Bartlett, S. E., & Bonci, A. (2011). Chlorzoxazone, an SK channel activator used in humans, reduces excessive alcohol intake in rats. Biological Psychiatry, 69(7), 618–624. https://doi.org/10.1016/j.biopsych.2010.11.011.Chlorzoxazone

Huang, L., Chen, Y., Jin, S., Lin, L., Duan, S., Si, K., Gong, W., & Julius Zhu, J. (2021). Organizational principles of amygdalar input-output neuronal circuits. Molecular Psychiatry, online ahe. https://doi.org/10.1038/s41380-021-01262-3

Janak, P. H., & Tye, K. M. (2015). From circuits to behaviour in the amygdala. Nature, 517(7534), 284–292. https://doi.org/10.1038/nature14188.From

Jaramillo, A. A., Williford, K. M., Marshall, C., Winder, D. G., & Centanni, S. W. (2020). BNST transient activity associates with approach behavior in a stressful environment and is modulated by the parabrachial nucleus. Neurobiology of Stress, 13, 100247. https://doi.org/10.1016/j.ynstr.2020.100247

Kash, T. L., Baucum, A. J., Conrad, K. L., Colbran, R. J., & Winder, D. G. (2009). Alcohol exposure alters NMDAR function in the bed nucleus of the stria terminalis. Neuropsychopharmacology, 34(11), 2420–2429. https://doi.org/10.1038/npp.2009.69.Alcohol

Keistler, C. R., Hammarlund, E., Barker, J. M., Bond, C. W., DiLeone, R. J., Pittenger, C., & Taylor, J. R. (2017). Regulation of alcohol extinction and cue-induced reinstatement by specific projections among medial prefrontal cortex, nucleus accumbens, and basolateral amygdala. Journal of Neuroscience, 37(17), 4462–4471. https://doi.org/10.1523/JNEUROSCI.3383-16.2017

Kim, S., Adhikari, A., Lee, S. Y., Marshel, J. H., Christina, K., Mallory, C. S., Lo, M., Pak, S., Mattis, J., Lim, B. K., Malenka, R. C., Warden, M. R., Neve, R., & Tye, K. M. (2013). Diverging neural pathways assemble a behavioural state from separable features in anxiety. Nature, 496(7444), 219–223. https://doi.org/10.1038/nature12018.Diverging

Koob, G. F., Buck, C. L., Cohen, A., Edwards, S., Park, P. E., Schlosburg, J. E., Schmeichel, B., Vendruscolo, L. F., Wade, C. L., Whit, T. W., & George, O. (2014). Addiction as a stress surfeit disorder. Neuropharmacology, 76, 370–382. https://doi.org/10.1016/j.neuropharm.2013.05.024

Koob, G. F., & Volkow, N. D. (2016). Neurobiology of addiction: a neurocircuitry analysis. The Lancet Psychiatry, 3(8), 760–773. https://doi.org/10.1016/S2215-0366(16)00104-8

Kuntsche, E., & Müller, S. (2011). Why do young people start drinking? Motives for first-time alcohol consumption and links to risky drinking in early adolescence. European Addiction Research, 18(1), 34–39. https://doi.org/10.1159/000333036

Läck, A. K., Ariwodola, O. J., Chappell, A. M., Weiner, J. L., & McCool, B. A. (2008). Ethanol inhibition of kainate recpetor-mediated excitatory neurotransmission in the rat basolateral nucleus of the amygdala. Neuropharmacology, 55(5), 661–668. https://doi.org/10.1016/j.neuropharm.2008.05.026.ETHANOL

Läck, A. K., Diaz, M. R., Chappell, A., DuBois, D. W., & McCool, B. A. (2007). Chronic ethanol and withdrawal differentially modulate pre-and postsynaptic function at glutamatergic synapses in rat basolateral amygdala. Journal of Neurophysiology, 98(6), 3185–3196. https://doi.org/10.1152/jn.00189.2007

Lange, M. D., Daldrup, T., Remmers, F., Szkudlarek, H. J., Lesting, J., Guggenhuber, S., Ruehle, S., Jüngling, K., Seidenbecher, T., Lutz, B., & Pape, H. C. (2017). Cannabinoid CB1 receptors in distinct circuits of the extended amygdala determine fear responsiveness to unpredictable threat. Molecular Psychiatry, 22(10), 1422–1430. https://doi.org/10.1038/mp.2016.156

Leichnetz, G. R., & Astruc, J. (1977). The course of some prefrontal corticofugals to the pallidum, substantia innominata, and amygdaloid complex in monkeys. Experimental Neurology, 54, 104–109. https://doi.org/10.1016/0014-4886(77)90238-2

Li, H., Penzo, M. A., Taniguchi, H., Kopec, C. D., Huang, Z. J., & Li, B. (2013). Experience-dependent modification of a central amygdala fear circuit. Nature Neuroscience, 16(3), 332–339. https://doi.org/10.1038/nn.3322.Experience-dependent

Marty, V. N., & Spigelman, I. (2012). Effects of alcohol on the membrane excitability and synaptic transmission of medium spiny neurons in the nucleus accumbens. Alcohol, 46(4), 317–327. https://doi.org/10.1016/j.alcohol.2011.12.002.Effects

McCool, B. A., Christian, D. T., Fetzer, J. A., & Chappell, A. M. (2014). Lateral/basolateral amygdala serotonin type-2 receptors modulate operant self-administration of a sweetened ethanol solution via inhibition of principal neuron activity. Frontiers in Integrative Neuroscience, 8, 1–13. https://doi.org/10.3389/fnint.2014.00005

McDonald, A. J. (1998). Cortical pathways to the mammalian amygdala. Progress in Neurobiology, 55, 257–332. https://doi.org/10.1016/S0301-0082(98)00003-3

McDonald, A. J., Mascagni, F., & Guo, L. (1996). Projections of the medial and lateral prefrontal cortices to the amygdala: A phaseolus vulgaris leucoagglutinin study in the rat. Neuroscience, 71(1), 55–75.

McGinnis, M. M., Parrish, B. C., Chappell, A. M., Alexander, N. J., & McCool, B. A. (2020). Chronic ethanol differentially modulates glutamate release from dorsal and ventral prefrontal cortical inputs onto rat basolateral amygdala principal neurons. ENeuro, 7(2), 1–17. https://doi.org/10.1523/ENEURO.0132-19.2019

McGinnis, M. M., Parrish, B. C., & McCool, B. A. (2020). Withdrawal from chronic ethanol exposure increases postsynaptic glutamate function of insular cortex projections to the rat basolateral amygdala. Neuropharmacology, 172, 108129. https://doi.org/10.1016/j.neuropharm.2020.108129

Millan, E. Z., Kim, H. A., & Janak, P. H. (2017). Optogenetic activation of amygdala projections to nucleus accumbens can arrest conditioned and unconditioned alcohol consummatory behavior. Physiology & Behavior, 360, 106–117. https://doi.org/10.1016/j.neuroscience.2017.07.044.Optogenetic

Morales, M., McGinnis, M. M., & McCool, B. A. (2015). Chronic ethanol exposure increases voluntary home cage intake in adult male, but not female, Long-Evans rats. Pharmacology, Biochemistry and Behavior, 139, 67–76. https://doi.org/10.1109/EMBC.2016.7590696.Upper

Morales, M., McGinnis, M. M., Robinson, S. L., Chappell, A. M., & McCool, B. A. (2018). Chronic intermittent ethanol exposure modulation of glutamatergic neurotransmission in rat lateral/basolateral amygdala is duration-, input-, and sex-dependent. Neuroscience, 371, 277–287. https://doi.org/10.1183/09031936.00063810

Namburi, P., Beyeler, A., Yorozu, S., Calhoon, G. G., Halbert, S. A., Wichman, R., Holden, S. S., Mertens, K. L., Anahtar, M., Felix-Ortiz, A. C., Wickersham, I. R., Gray, J. M., & Tye, K. M. (2015). A circuit mechanism for differentiating positive and negative associations. Nature, 50(7549), 675–678. https://doi.org/10.1126/science.1249098.Sleep

Oliver, D. K., Intson, K., Sargin, D., Power, S. K., McNabb, J., Ramsey, A. J., & Lambe, E. K. (2020). Chronic social isolation exerts opposing sex-specific consequences on serotonin neuronal excitability and behaviour. Neuropharmacology, 168, 108015. https://doi.org/10.1016/j.neuropharm.2020.108015

Overstreet, D. H., Knapp, D. J., & Breese, G. R. (2004). Similar anxiety-like responses in male and female rats exposed to repeated withdrawals from ethanol. Pharmacology, Biochemistry and Behavior, 78(3), 459–464. https://doi.org/10.1016/j.pbb.2004.04.018.Similar

Padula, A. E., Griffin, W. C., Lopez, M. F., Nimitvilai, S., Cannady, R., McGuier, N. S., Chesler, E. J., Miles, M. F., Williams, R. W., Randall, P. K., Woodward, J. J., Becker, H. C., & Mulholland, P. J. (2015). KCNN genes that encode small-conductance Ca 2+-activated K+ channels influence alcohol and drug addiction. Neuropsychopharmacology, 40(8), 1928–1939. https://doi.org/10.1038/npp.2015.42

Pati, D., Marcinkiewcz, C. A., DiBerto, J. F., Cogan, E. S., McElligot, Z. A., & Kash, T. L. (2020). Chronic intermittent ethanol exposure dysregulates a GABAergic microcircuit in the bed nucleus of the stria terminalis. Neuropharmacology, 168, 107759. https://doi.org/10.1016/j.neuropharm.2019.107759.Chronic

Pleil, K. E., Helms, C. M., Sobus, J. R., Daunais, J. B., Grant, K. A., & Kash, T. L. (2016). Effects of chronic alcohol consumption on neuronal function in the non-human primate BNST. Addiction Biology, 21(6), 1151–1167. https://doi.org/10.1111/adb.12289.Effects

Pleil, K. E., Lowery-gionta, E. G., Crowley, N. A., Li, C., Catherine, A., Rose, J. H., Mccall, N. M., Maldonado-devincci, A. M., Morrow, L., Jones, S. R., & Kash, T. L. (2015). Effects of chronic ethanol exposure on neuronal function in the prefrontal cortex and extended amygdala. Neuropharmacology, 99, 735–749. https://doi.org/10.1016/j.neuropharm.2015.06.017.Effects

Rau, A. R., Chappell, A. M., Butler, T. R., Ariwodola, O. J., & Weiner, J. L. (2015). Increased basolateral amygdala pyramidal cell excitability may contribute to the anxiogenic phenotype induced by chronic early-life stress. Journal of Neuroscience, 35(26), 9730–9740. https://doi.org/10.1523/JNEUROSCI.0384-15.2015

Ray, M. H., Russ, A. N., Walker, R. A., & McDannald, M. A. (2020). The Nucleus Accumbens Core is Necessary to Scale Fear to Degree of Threat. Journal of Neuroscience, 40(24), 4750–4760. https://doi.org/10.1523/JNEUROSCI.0299-20.2020

Sah, P., Faber, E. S. L., Lopez De Armentia, M., & Power, J. (2003). The amygdaloid complex: anatomy and physiology. Physiological Reviews, 83(3), 803–834. https://doi.org/10.1152/physrev.00002.2003

Schulteis, G., Markou, A., Cole, M., & Koob, G. F. (1995). Decreased brain reward produced by ethanol withdrawal. Proceedings of the National Academy of Sciences of the United States of America, 92, 5880–5884. https://doi.org/10.1073/pnas.92.13.5880

Scott, H., Tjernström, N., & Roman, E. (2020). Effects of pair housing on voluntary alcohol intake in male and female Wistar rats. Alcohol, 86, 121–128. https://doi.org/10.1016/j.alcohol.2019.12.005

Senn, V., Wolff, S. B. E., Herry, C., Grenier, F., Ehrlich, I., Gründemann, J., Fadok, J. P., Müller, C., Letzkus, J. J., & Lüthi, A. (2014). Long-range connectivity defines behavioral specificity of amygdala neurons. Neuron, 81, 428–437. https://doi.org/10.1016/j.neuron.2013.11.006

Shan, L., Galaj, E., & Ma, Y. Y. (2019). Nucleus accumbens shell small conductance potassium channels underlie adolescent ethanol exposure-induced anxiety. Neuropsychopharmacology, 44(11), 1886–1895. https://doi.org/10.1038/s41386-019-0415-7

Sizer, S. E., Parrish, B. C., & McCool, B. A. (2021). Chronic Ethanol Exposure Potentiates Cholinergic Neurotransmission in the Basolateral Amygdala. Neuroscience, 455, 165–176. https://doi.org/10.1016/j.neuroscience.2020.12.014

Stuber, G. D., Sparta, D. R., Stamatakis, A. M., van Leeuwen, W. A., Hardjoprajitno, J. E., Cho, S., Tye, K. M., Kempadoo, K. A., Zhang, F., Deisseroth, K., & Bonci, A. (2011). Excitatory transmission from the amygdala to nucleus accumbens facilitates reward seeking. Nature, 475(7356), 377–380. https://doi.org/10.1038/nature10194

Substance Abuse and Mental Health Services Administration. (2019). Results from the 2018 National Survey on Drug Use and Health: Detailed tables. https://www.samhsa.gov/data/

Sugiyama, A., Yamada, M., Saitoh, A., Nagase, H., Oka, J. I., & Yamada, M. (2018). Administration of a delta opioid receptor agonist KNT-127 to the basolateral amygdala has robust anxiolytic-like effects in rats. Psychopharmacology, 235, 2947–2955. https://doi.org/10.1007/s00213-018-4984-7

Sun, P., Zhang, Q., Zhang, Y., Wang, F., Wang, L., Yamamoto, R., Sugai, T., & Kato, N. (2015). Fear conditioning suppresses large-conductance calcium-activated potassium channels in lateral amygdala neurons. Physiology and Behavior, 138, 279–284. https://doi.org/10.1016/j.physbeh.2014.10.005

Tye, K. M., Prakash, R., Kim, S., Fenno, L. E., Grosenick, L., Zarabi, H., Thompson, K. R., Gradinaru, V., Ramakrishnan, C., & Deisseroth, K. (2011). Amygdala circuitry mediating reversible and bidirectional control of anxiety. Nature, 471(7338), 358–362. https://doi.org/10.1038/nature09820.Amygdala

Vetter-O’Hagen, C. S., & Spear, L. P. (2011). The effects of gonadectomy on age-and sex-typical patterns of ethanol consumption in Sprague-Dawley rats. Alcoholism: Clinical and Experimental Research, 35(11), 2039–2049. https://doi.org/10.1111/j.1530-0277.2011.01555.x

Walker, D. L., & Davis, M. (1997). Double dissociation between the involvement of the bed nucleus of the stria terminalis and the central nucleus of the amygdala in startle increases produced by conditioned versus unconditioned fear. Journal of Neuroscience, 17(23), 9375–9383. https://doi.org/10.1523/jneurosci.17-23-09375.1997

Yin, F., Guo, H., Cui, J., Shi, Y., Su, R., Xie, Q., Chang, J., Wang, Y., & Lai, J. (2019). The basolateral amygdala regulation of complex cognitive behaviours in the five-choice serial reaction time task. Psychopharmacology, 236, 3135–3146. https://doi.org/10.1007/s00213-019-05260-w

